# Swarming bacterial fronts: Dynamics and morphology of active swarm interfaces propagating through passive frictional domains

**DOI:** 10.1101/2020.04.18.048637

**Authors:** Joshua Tamayo, Yuchen Zhang, Merrill E Asp, Alison E Patteson, Arezoo M Ardekani, Arvind Gopinath

**Affiliations:** Department of Bioengineering, University of California Merced, Merced, CA; Department of Mechanical Engineering, Purdue University, Lafayette, IN; Department of Physics, Syracuse University, Syracuse, NY

## Abstract

Swarming, a multicellular mode of flagella-based motility observed in many bacteria species, enables coordinated and rapid surface translocation, expansion and colonization. In the swarming state, bacterial films display several characteristics of active matter including intense and persistent long-ranged flocks and strong fluctuating velocity fields with significant vorticity. Swarm fronts are typically dynamically evolving interfaces. Many of these fronts separate motile active domains from passive frictional regions comprised of dead or non-motile bacteria. Here, we study the dynamics and structural features of a model active-passive interface in swarming *Serratia marcescens*. We expose localized regions of the swarm to high intensity wide-spectrum light thereby creating large domains of tightly packed immotile bacteria. When the light source is turned off, swarming bacteria outside this passivated region advance into this highly frictional domain and continuously reshape the interphase boundary. Combining results from Particle Image Velocimetry (PIV) and intensity based image analysis, we find that the evolving interface has quantifiable and defined roughness. Correlations between spatially separated surface fluctuations and damping of the same are influenced by the interaction of the interphase region with adjacently located and emergent collective flows. Dynamical growth exponents characterizing the spatiotemporal features of the surface are extracted and are found to differ from classically expected values for passive growth or erosion. To isolate the effects of hydrodynamic interactions generated by collective flows and that arising from steric interactions, we propose and analyze agent-based simulations with full hydrodynamics of rod-shaped, self-propelled particles. Our computations capture qualitative features of the swarm and predict correlation lengths consistent with experiments. We conclude that hydrodynamic and steric interactions enable different modes of surface dynamics, morphology and thus front invasion.

## 1 Introduction

Swarming migration or swarming in short is a multicellular mode of flagella-based motility ^1–4^ featuring collective long-ranged flows that allows bacterial colonies to rapidly and efficiently cover and subsequently colonize surfaces ^5–12^. The swarming response can be initiated by transferring planktonic free-swimming bacteria in liquid media to soft agar gels. This process results in a change to bacterial phenotypes that are morphologically different (longer with more flagella) ^5,9^.

Swarming bacterial films share several characteristics, hyper-flagellated and usually elongated cells, secretion of a wetting agent, and a high-density population of specialized swarmers ^7,8^ all of which drive cooperative movement. In terms of structural features, swarming films are usually a monolayer a few millimeters wide near the leading edge of the swarm; far from this boundary, multilayered growth is observed ^1–4^. As in biofilm formation ^13–16^ (that is distinct and develops differently from swarming), swarms are dense and characterized by strong inter-cell interactions. Characteristics of the state as well as onset of swarming depends strongly on the nutrient composition of the liquid in the substrate as well as viscosity of the ambient medium. Unlike biofilms, however, the motile swarms feature long-range collective motions ^7,9,17–25^ featuring structures with defined length scales and persistence times.

Our recent studies ^24,25^ showed that the flow features respond to the geometry as well as the frictional resistance of the environment in which the swarms move. From a biological standpoint, these emergent flow characteristics thus seem to confer benefits relevant to colonization and occupation of preferred niches ^6,7,26,27^. Additionally, enhanced surface expansion rates may help fitness ^6^ and yield elevated resistance to antibiotics ^27^. Swarming may thus contribute to the ability of pathogenic species to cause and spread infections, in part due to enhanced motility and in part due to the co-regulation of associated virulence factors.

Besides being relevant in biology and medicine, swarming bacteria are a convenient experimental system to study and test models for dense active matter and active fluids ^23,28–32^. Recent numerical and theoretical studies have focused on the collective flows and identified dynamical exponents characterizing the energy spectrum and spatiotemporal features as well as variations near interfaces ^24,28^. To explain these features, theoretical models for swarming systems based on adaptations of classical nematic hydrodynamic theories and hydrodynamic multiphase models have been used to characterize the phase-separating active nematic and passive phases and propagation of interfaces of active nematics on substrates ^33–36^. Understanding the role of swarming in enabling bacterial collective motility and rapid propagation is both important and timely; this will require combination of experimental, analytical and computational approaches.

An important feature of bacterial swarms are the dynamically evolving interfaces and propagating fronts that form naturally. A prototypical example of these fronts are the interfaces that separate motile active domains from passive highly frictional regions comprised of dead or non-motile bacteria. In recent work ^24,25^, we used the response to wide-spectrum light to create internal active-passive interfaces within dense active swarms of *Serratia marcescens*. The overall effect of light was found to depend on both exposure time and intensity; high intensities coupled with long exposure times caused a permanent loss of motility ^25^. Combined with suitable screens, this provided a controllable method to generate macroscopic domains with nearly circular or straight interfaces. The passive domain and active swarm interact at this interphase boundary where self-emergent, vortical flows develop ^24^. Our experiments revealed that the interface region has a well-defined thickness, with a quantifiable surface energy and raise the possibility that the propagation speed of the diffuse interface couples to local swarm velocity and interface curvature ^24,37^.

In this article, we examine and investigate an equally important aspect of the active-passive interface, namely its spatiotemporal structure that enables the manner of propagation and colonization. The passive region comprises of densely packed aligned bacteria arranged in static domains that are typically smaller than the structure of the collective flow structures. Over length scales associated with multiple domains, the passive region appears as a highly frictional region which the active bacteria have to penetrate in order to propagate. Employing PIV techniques and intensity based image analysis, we find that the diffuse, moving interphase region is characterized by a well-defined roughness profile. Correlations between spatially and temporally separated surface undulations are controlled by the interaction of the interface region with the intense adjacently located active flows. Dynamical and growth exponents characterizing the spatiotemporal roughness profile differ from the canonical values valid for passive growth or erosion. Our experiments also suggest possible self-similar behavior for the experimentally measured interfacial roughness opening up a question for future exploration.

It is further of interest to understand if steric effects and hydrodynamic effects play equal roles in the interface propagation and in determining the spatiotemporal features of the interface, specifically roughness features and swarming intensity. Furthermore, understanding the relative importance of hydrodynamic interactions will clarify the role of fluid content in the underlying substrate. Our experiments on *Serratia marcescens* do not allow us to discriminate and separate the impact of purely steric effects from fluid mediated hydrodynamic interactions on the collective motion well within the active phase as well as in the vicinity of the interface. To investigate each effect separately and also their combined effect when acting in tandem, we analyze a discrete agent based simulations that treat the bacteria as inertialess self-propelled rods (SPR) moving in a viscous fluid in two dimensions with parameters chosen to mimic the bacterial system.

The paper is organized as follows. In Section 2, we present experimental results and analysis of active swarming bacteria moving into a densely packed domain of immotile bacteria. We focus on analysis of interfacial structure and morphology and roughness, complementing our previous studies on the speed of propagation ^24,25^. Details of the agent based simulation model, scheme and implementation details are presented first in Section 3, followed by a discussion of our results in Section 4. We first compare features of the simulated swarm with experimental data to check for consistency and fidelity. We then analyze results for the simulated swarms without and then with hydrodynamic interactions and study the changes induced in interface morphology and invasion dynamics. Taken together, our experiments and simulations suggest that hydrodynamic and steric interactions enable different modes of surface dynamics, morphology and front invasion. We conclude in Section 5 by summarizing our results and suggesting avenues for further experimental and theoretical exploration.

## 2 Experiments: Swarming *Serratia marcescens* as a model dense active system

We use swarming *Serratia marcescens* as our model system to study two aspects of the swarming process as an active swarm propagates into a passive medium offering significant resistance - 1) the spatiotemporal structure of the interface and changes in morphology (distinct from the mean speed analyzed in ^24^as it propagates, and 2) understanding the interplay of direct (steric) cell-cell interactions, emergent active flows such as streamers, flocks and vortices generated in the interphase region and fluid mediated cell-cell interactions.

The experimental protocols to prepare the swarms of *Serratia marcescens* and the method of creating immotile (passive) domains within actively swarming regions have been detailed previously ^24,25^. A brief summary that highlights aspects important to the current work is provided in Appendix A at the end.

### 2.1 Results

#### 2.1.1 Features of the swarming base state prior to exposure

As a point of departure we summarize observations of our base state - the unexposed actively swarming region. In collectively-moving swarms, individual self-propelling cells are influenced by steric and hydrodynamic interactions with their neighbors ^7,9,10,19,24^. These interactions result in complex structural and flow features including fluctuating regions of high vorticity and streamers as seen in Figure 1(a) (see also ^24,25^) that superimposes PIV derived velocity fields on a static image of the active swarm.The intensity and transient of these emergent flows increases as one moves from the leading front of the swarm as indicated by the faint white curve in Figure 1(a).

**Fig. 1.**
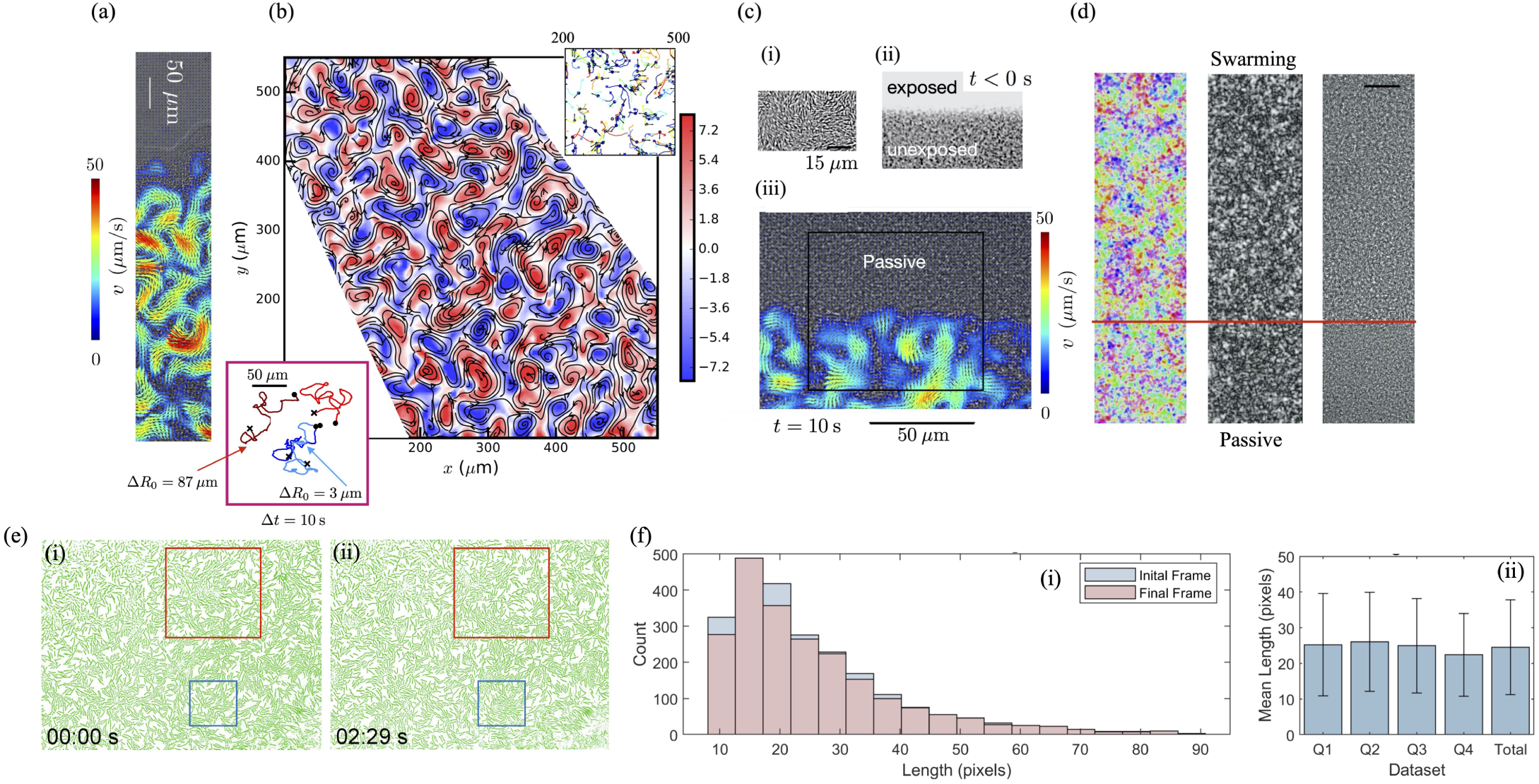
(a) (left) A section of a swarm front with velocity fields using PIV superimposed ^38,39^. The direction of propagation is upward. The free edge of the swarm is seen on the top as the white curve. Intensity of the swarm increases rapidly as one moves into the swarm. The substrate is agar. We note that swarm speeds can reach values of 50 *µ*m/s. (right) Snapshot of the vorticity field in the active phase that illustrates the alternate arrangement of well defined vortical structures. (b) (Inset, Top right) The vortical and time-dependent nature of the collective flows are easily observed by seeding the swarm film with 2*µ*m particles (top, scale bar 50 *µ*m) and then tracking their absolute as well as relative motions over time (Inset, Bottom left). (Center) We note persistent motion for small times leading to complex diffusion-like response. Tracers that are close to each other may move apart rather rapidly. The length scales of the vortical structures are ∼ 20 − 25 *µ*m; the residence time of structures is around 0.8 seconds ^24^. Also note the presence of vortices of both signs as indicated by the organized arrangement of blue and red colored areas. Note the vortex lengths comprise of a few bacterial lengths. (c) (i) Raw image of the active (dark) - passive (light) edge generated by light exposure. (ii) Examination of the vicinity of the front separating the active and passive domains 10s after exposure (*t* > 0) reveals strong velocity gradients (in color) in the active region. In the passive phase, immotile bacteria reside in jammed states until they are swept away by the advancing front. Far from the interface, the active flows show intense vortical patterns, with clockwise and counter-clockwise patterns arrayed periodically. These vortices are transient and constantly evolving and are interspersed with other features such as streamers and flocks. Snapshot from video taken with 63x (NA = 0.7) objective in a Nikon inverted microscope and includes a larger area of view. (d) Structural analysis using *ImageJ* and the *OrientationJ* plugin of a section of the active-passive interface. Snapshot from raw video acquired using 20x Nikon objective at 30 fps. Passivated bacteria are to the bottom, the active swarm erodes this passive region as it moves from top to bottom. We show (left) - length scales of similarly clusters, (middle, grayscale indicating degree of alignment) the degree of alignment and (right) an instantaneous snapshot. (e) (i, ii) Snapshots at two different times in the low density non-swarming state nonetheless provide significant insight into how bacterial clusters navigate jammed, stuck or otherwise immotile bacteria. ESM Movie 1 provides clear evidence of the manner in which steric interactions align flocks and clusters leading to efficient displacement in frictional regions with steric obstacles. Images are thresholded and color contrasted snapshots from raw videos. (f) Analysis using the *Directional J* plugin of the length distributions corresponding to the first and last frames in ESM Movie 1. Here the length distributions (average length ∼ 8.1 *µ*m) overlap suggesting the tracking algorithms we use are accurate. The wide variation in length shows how the bacteria - even prior to onset of swarming - can attain varying lengths and move differently.

The probability distribution function *p*(*v*) of averaged speeds (based on the PIV-measurements) *v* in a region of area 400 *µ*m^2^ and at a distance 100 *µ*m from the edge of the expanding colony shows a peak at 18 *µ*m/s, an expectation value ≈ 28 *µ*m/s and a tail that extends to 100 *µ*m/s. The advancing front of the swarm is approximately 3 *µ*m/s ^24^. The spatial correlation function *C*_*v*_(Δ*r*) and the temporal correlation *C*_*t*_ (Δ*t*) of the velocity fields were also calculated using

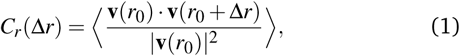

and

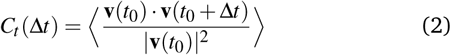

respectively, with brackets denoting averages over reference positions *r*_0_ and times *t*_0_. Application of equation (1) yielded typical vortex sizes *λ*_c_ ≈ 20 *µ*m. For Δ*r* < *λ*_c_, bacterial velocities are correlated positively, *C*_*r*_ > 0. As Δ*r* increases, the function *C*_*r*_ decays crossing zero and then stays negative for 21 *µ*m < Δ*r* < 32 *µ*m indicating neighboring vortices are typically anti-correlated. For Δ*r* > 32 *µ*m, velocity fields become progressively uncorrelated. Fitting *C*_*t*_ (Δ*t*) = exp(−Δ*t/τ*), we estimate *τ* ≈ 0.25 s.

In Figure 1(b), we show a snapshot of the PIV derived vorticity field in the interior of the swarm region away from any interfaces. Note the organized, arrayed groups of clockwise and counter-clockwise vortices. The intensity and temporary nature of these vortices is clearly evidenced upon seeding the swarm with small 2 *µ*m colloidal polystyrene spheres; these particles are strongly advected by the bacteria and follow tortuous trajectories (upper right, Figure 1(b)). The particle trajectories obtained from standard particle tracking techniques are shown for 4 second time intervals. We also determined the particle speed distribution *p*(*v*) by pooling together the particle speed over time for hundreds of particles; the particle speed is defined as the two-dimensional particle displacement over a 1 second time interval. The particle speed distribution measurement (blue circles) was found to follow a 2D Maxwell-Boltmann distribution ^24,25^, *p*(*v*) = *vm*(*k*_*B*_*T*_eff_)^−1^exp(−*mv*^2^*/*2*k*_*B*_*T*_eff_), where *m* is the mass of the polystyrene particle, *k*_*B*_ is the Boltzmann constant, and *T*_eff_ ≈ 2.2 × 10^5^ K, approximately 700 times the thermal temperature (293 K).

The effective temperature may be interpreted as a mixture temperature purely due to the energy in the swarming collective flows. The mixing produced by these fluctuating flows is clearly illustrated in the inset below (bottom left, Figure 1(b)) where we plot trajectories of spatially separated tracers (initial separations 3 *µ*m and 87 *µ*m) tracers. The mean square displacement obtained from long time measurements provides an effective diffusivity that characterizes the activity of the suspension.

#### 2.1.2 Collective flows drive the propagating front

We next examine the collective velocity fields around the inter-phase region that straddles the active and passive phases. Figures 1(c) (i)-(ii) illustrate the interphase region of the swarm (shown magnified in (i)) and also during light exposure (Figure 1 (c) - (ii)). The top half of the image (bright) is the domain that is exposed to light incident through a half-plane aperture. The dark (bottom) half is the unexposed active swarm. We note the (apparent) boundary between the two phases is straight on scales much larger than the length scale of the correlated structures but diffuse and mushy on shorter length scales.

Focussing on the interphase region in Figure 1(c)-(iii),we show the velocity fields obtained from PIV analysis superposed on the raw image. Swarming bacteria in the active segment of the figure exhibit strong collective motion *right up to* the boundary region (flow speeds ∼ 50 *µ*m/s). Vortices align along the interface, and are arrayed normal to it as well. These dynamic vortices (typical lifetime, ∼ 0.24 s, frequency 0.8 s, characteristic vortex size ∼ 20 − 25 *µ*m) etch the interface into cusps and valleys and are observed to control the morphology and structure of the interface region.

Theoretical and computational studies of dense suspensions of passive rods suggest that polar and/or nematic alignment may be induced due to a combination of excluded volume effects (modeled for instance using a mean-field Maier-Saupe or Onsager potential) and thermal diffusion ^40,41^. Recognizing that bacterial swarms may behave similarly, we examined the structure and alignment of bacterial clusters in the active and passive phases as the interface position changed in time using the *Directional J* and *OrientationJ* plugins that are a part of the open source software ^38^ *ImageJ*\* Qualitatively, the orientation is visualized as a false-color image with colored domains with domains of the same color indicating similarly aligned bacteria and intensity quantifying degree of coherence. Structural information such as size of coherently arrayed structures and length scales over which structural features are correlated may thus be obtained. Using these plugins we found that these domains exhibited a distribution of sizes, typically between 10-20 *µ*m with continuously changing orientational fields due to emergent flows ^25^. Figure 1(d) shows the result of such an image based analysis of the structural features demonstrating the presence of many intertwined aligned domains. This however is just a static representation of the highly dynamical swarming process.

#### 2.1.3 Intensity field quantify interface shape and roughness

The raw videos obtained from the images provide a sequence of snapshots of the intensity field over the domain of observation as a function of time. Noting that the the mean intensity fluctuations in the active and passive phases far from the boundary, |Δ*I*_*A*_(*t*)| and |Δ*I*_*P*_(*t*)|, remained relatively constant over the duration of the longest experiments, we defined a scalar order parameter, *ϕ*_E_ ∈ [−1, 1] (the subscript E denotes experiments)

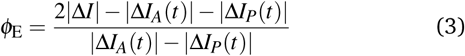

that was computed from intensity fluctuations Δ*I* between successive images to locate and track the inter-phase boundary. To reduce noise in the system (due to pixel resolution, short-range fluctuations and background light fluctuations), we then filtered the pixel-wise calculated order parameter by smoothing the data over a 3 × 3 *µ*m^2^ area. The filtered phase field has value −1 corresponding to the completely passive phase and +1 corresponding to the fully active phase. The interphase boundary was identified as the characteristic region over which *ϕ*_E_ changes from one asymptotic value to the other; sharp variation occurring over a finite length that is identified ^24^ as a characteristic interface width.

To obtain a clear value for the mean interface position, we therefore defined the interface position (in the two dimensional *x* − *y* plane) **r**_Int_(*t*) implicitly by the locus of points such that *ϕ*_E_(**r**_Int_, *t*) = 0. Our system being approximately two-dimensional, we chose to orient the *x* axis along the edge of active-passive interface observed at the instant exposure is stopped *t* = 0. The interface shapes extracted from the phase field did not feature overhangs.

Time snaps of the interface location (here the two dimensional curve in the *x* − *y* plane) for two representative experiments are shown in Fig, 2(a) - (i) & (ii) along with a schematic sketch of the notation and definitions used. We note two main features characterizing the spatiotemporal evolution of the interface profiles. First, the profiles clearly exhibit significant roughness. Second, the interface propagates due to the active domain eroding the passive one. In Figure 2(b), we show the PIV derived snapshots of the vorticity field and overlaid streamlines for the experiment corresponding to Figure 1(c)-(iii) illustrating the rapid turnover of collective flow features before the interface has the time to propagate significantly. We observe the dynamic motion of individual vortices that cause the interface (solid blue curve) to propagate and roughen. Individual vortices - red (counter-clockwise) and blue (clockwise) - etch the interphase boundary (blue line).

**Fig. 2.**
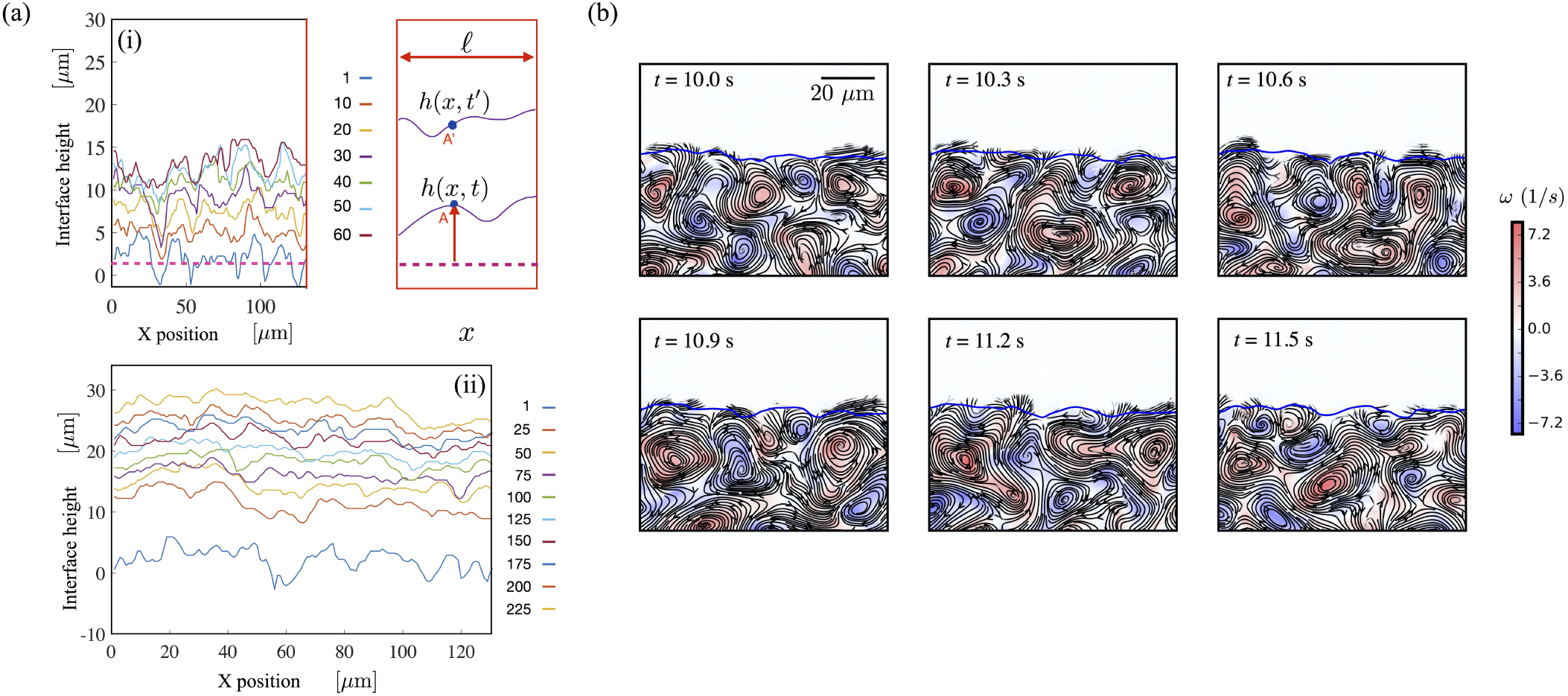
(a) Interface profiles in the (*x, y*) plane obtained from the phase-field based on intensity fluctuations for two experimental data sets - (i) & (ii). The shapes are obtained by fitting a vertically averaged (along *y*) one-dimensional phase field and seeking the location of *ϕ*_E_ = 0 as a function of the lateral position, *x*. Time stamps are in seconds. For these profiles, the one-dimensional, averaged phase field (system size *L* = 200 *µ*m) 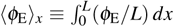 follows a quasi-static form corresponding to a diffuse interface that slowly propagates with finite thickness. Also shown is a schematic demonstrating the definitions of the variables used to calculate the interface location and thence the interface height *h*(*x, t*). (b) Snapshots focusing on a subdomain of Figure 1(b). The filtered two-dimensional interface position identified by the locus *ϕ*_E_(**r**_Int_, *t*) = 0 is shown as the blue curve for various times after cessation of exposure. PIV derived Snapshots of the vorticity field and overlaid streamlines reveal the dynamic motion of individual vortices that cause the interface (black curve) to propagate and roughen. Individual vortices - red (counter-clockwise) and blue (clockwise) - etch the interphase boundary (blue line).

Averaging over the lateral dimension normal to the direction of propagation (c.f profiles in Figure 2(a), direction of arrow in the schematic, maximum overall system size *L* = 200 *µ*m), we obtain a one-dimensional, phase field profile 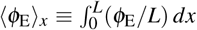 that, after initial transients, is well represented by

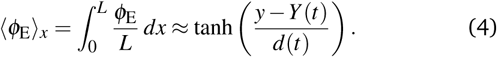

Equation (4) provides the *mean averaged* interface location *Y*(*t*) and the intrinsic thickness *d*(*t*). Details of the procedure and the connection between the spatiotemporal variations in *d* and *Y* and the collective flow in the active phase have been analyzed ^24,25^ in previous studies and will not be addressed here.

#### 2.1.4 Spatio-temporal correlations and roughness profiles

Interface roughness is related to the extrinsic thickness *W*(the width correlation function) ^42–45^, that is defined as

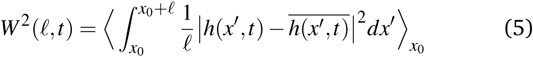

where 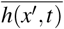 is the local average over distance *ℓ, x*_0_ is the reference position and *ℓ* the averaging box length that in our series of experiments varies from around 2 *µ*m up to the system size *L* = 200 *µ*m.

Figure 3 shows sample interface roughness profiles *W*(*ℓ, t*): Focusing first on the *ℓ* dependence of *W* - shown in Fig 3(a) – we observe two regimes, the first corresponding to *ℓ* < 80 *µ*m and the second to *ℓ* > 80 *µ*m. For either regime, *W* increases with *t*(blue to red) with the curves eventually collapsing to a single curve (c.f the orange at 53 s and the red curve at 66 s). A time average of the width *W* is shown in fig. 3(b) - also shown is the experiment to experiment variation (grey region, see caption). We find ⟨*W*⟩_*t*_ ∼ *ℓ* for *ℓ* < 80 *µ*m. For *ℓ* > 80 *µ*m, the dependence is very weak (the exponent being smaller than 1/2). Together, the data in Figs. 3(a) & 3(b) suggest that for interfacial separations less than ≈ 80 *µ*m, roughness is perhaps correlated (power law > 1*/*2). This result may be a consequence of the periodicity of the active vortical flows combined with our observation that the surface may exhibit resistance to deformation. However a clear understanding of the length scale is elusive.

**Fig. 3.**
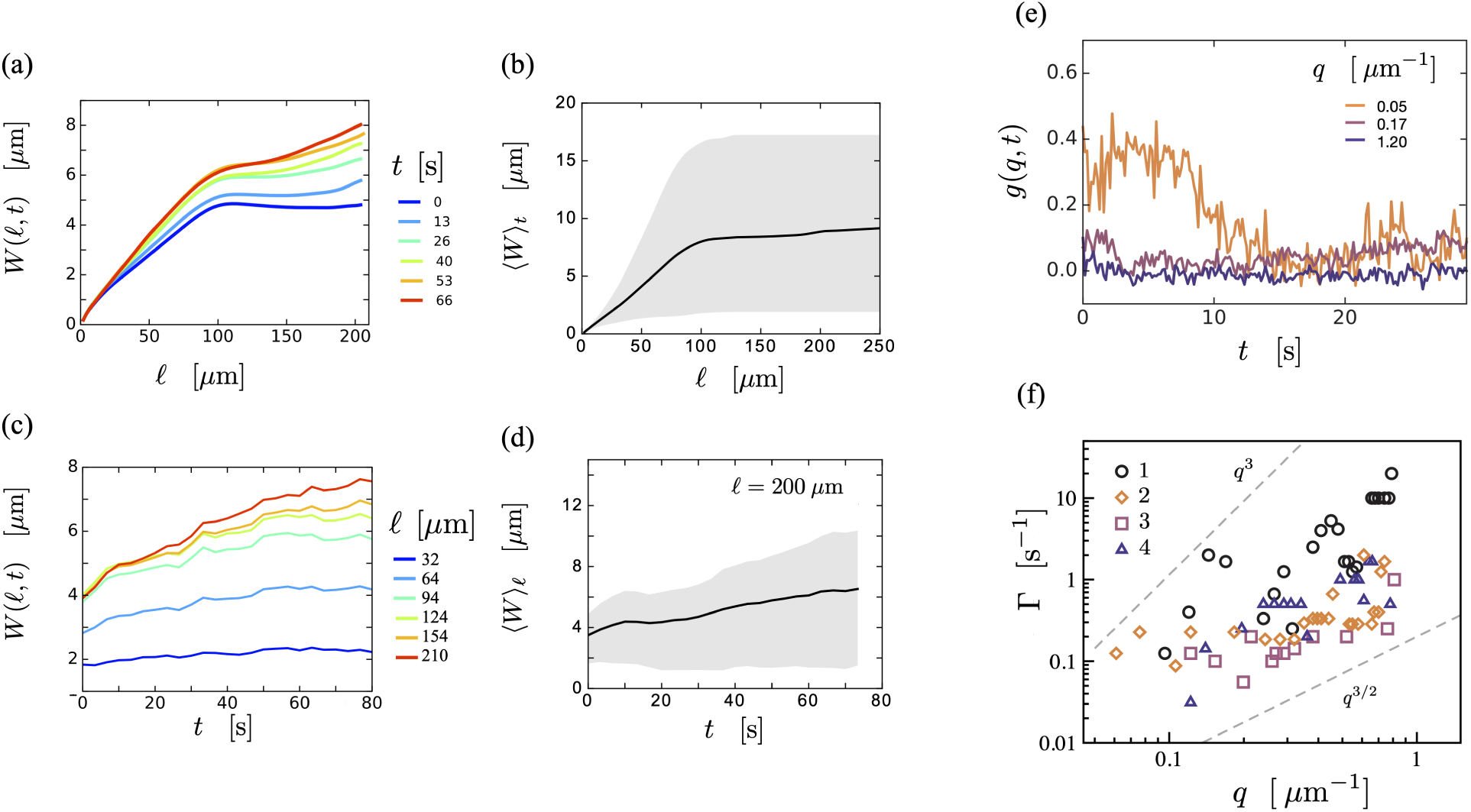
Interface roughness as a function of box length and time and its statistical properties. The upper and lower roughness bounds, are measured from four experiments. (a) Local interface roughness profiles *W* as a function of box length *ℓ* measured over time *t* for a single experiment. (b) Averaging over time, the mean interface roughness ⟨*W*⟩_*t*_ versus *ℓ* increases sharply for *ℓ* < 80 *µ*m, above which the roughness remains approximately constant. We find ⟨*W*⟩_*t*_ ∼ *ℓ* for *ℓ* < 80 *µ*m. For *ℓ* > 80 *µ*m up to the system size *L*, the dependence is very weak. The grey area show upper and lower limits from four experiments indicating the sample to sample variation from the mean. We find that ⟨*W*⟩_*t*_ ∼ *ℓ* for *ℓ* < 80 and scales weaker than 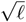 for *ℓ* > 80. The mean value corresponds to approximately the length of a swarming bacterial cell (5-10 *µ*m) and is much smaller than the scale of the vortical flows adjacent to the interface (20 *µ*m). (c) Sample interface roughness *W* values for various *ℓ* in a single experiment as a function of time, *t*. (d) Averaging over multiple experiments (with system size, *L* = 200 *µ*m held fixed), *W* increases from approximately 4 to 6 *µ*m over 70 seconds. We find ⟨*W*⟩ to be nearly constant for *t* < 25 s and then scale weakly with time (the power law exponent being between 1/2 and 1/3) for *t* > 25 s. (e) The correlation function 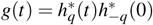 decays over time *t* for varying wavenumber *q*. We see oscillations that are superimposed on the mean decay suggesting that fluctuations originating from the transient nature of the near-interface active flow impact the damping rate. (f) Damping rates Γ(*q*) extracted from four experiments are clustered such that Γ ∼ *q*^*z*^, where 3*/*2 < *z* < 3.

We use our experimental data to analyze the interface roughness *W* values for various *ℓ* in a single flat interface experiment as a function of time, *t*. As seen in Fig. 3(c), we observe an increasing power law in time with increasing *ℓ*; of course, eventually for large *ℓ*(up to *ℓ* = *L*), we observe the curves approaching each other. Erosion is caused by the swirling vortical flows that form (arrayed periodically) adjacent to the interface. This spatial organization combined with time dependence of the positions at which vortices are located, results in fluctuations in *W* observed in Fig 3(c). Fixing *ℓ* = 200 *µ*m, we average over multiple experiments and observe that the interface roughness varies very little with time for the first 25 seconds or so (Fig 3(d); note also that vortex lifetimes are < 1 s), eventually after this grows very weakly.

#### 2.1.5 Damping of interface fluctuations

Treating the interface as a continuous line, the global (system scale) structure factor *S*_*q*_ for wavenumber *q* = *nπ/L* may be related to the Fourier amplitudes 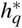 by 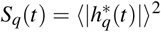. This and the wavenumber dependent autocorrelation function *g*

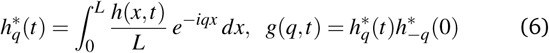

may then be easily evaluated. For a damped interface that fluctuates in response to external noisy forcing, fitting the growth coefficients to exp(−Γ(*q*)Δ*t*) cos(*ω*(*q*)Δ*t*) can help identifies regimes for which the damping is strong. Figs. 3(e) and 3(f) suggest that the interface responds in a complex manner with fluctuations that depend on wavenumber *q*. Note that these oscillations arise from a combination of passive thermal noise (thermodynamic) as well as additional non-equilibrium agitations due to the swirling, fluctuating flows at the interface. It is difficult to separate these two contributions from the data.

A passive fluid-fluid interface excited by thermal capillary waves may be treated as an harmonic oscillator driven by random noise that satisfies the fluctuation-dissipation theorem. Assuming a non-Markovian general Langevin equation with random forces that satisfy fluctuation-dissipation, Gross and Varnik ^46^ have recently derived scalings for the oscillation frequency and the growth rate in terms of density *ρ*, surface tension *σ*, and viscosity *µ*. In the weakly damped limit, they find *ω* ∼ *q*^3*/*2^ and Γ ∼ *q*^2^. In the strongly damped limit, they find Γ ∼ *q*.

Our extracted growth rates seems to suggest that the interface dampening falls in between these two limiting regimes; however variations between experiments make it difficult to extract the frequencies *ω* clearly. The dynamic growth exponent *z* defined through Γ(*q*) ∼ *q*^*z*^ are shown in Fig. 3(f); we find that Γ ∼ *q*^*z*^, where 3*/*2 < *z* < 3 with the damping rate increasing with *q*.

#### 2.1.6 Seeking self-similar roughness profiles

Dynamic scaling approaches for the description of initially flat, self-affine, single-valued interfaces that roughen with time have been proposed based on continuum models of the growth/erosion process in a variety of geophysical and materials physics contexts ^42,43,45,48^. Self-similar roughness profiles in these theoretically studied cases follow scalings given by *W*(*L, t*) ∼ *t*^*β*^, (*t* ≪ *t*^*^), *W*(*L, t*) ∼ *ℓ*^*α*^, (*t» t*^*^) and *t*^*^ ∼ *ℓ*^*z*^ with *z* = *α/β*. Here *α, β* and *ℓ* are the local roughness exponent, local growth exponent and system size respectively. The scaling relation here means that for early time (*t* ≪ *t*^*^), the roughness grows as a power law of time till it saturates for characteristic time *t*^*^. After time *t*^*^, the roughness grows as a power law of *ℓ* with *z* defined as the dynamic exponent,

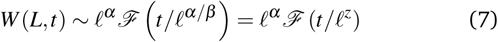

A quantity complementary to characteristic time *t*^*^, (if it exists) is the characteristic width *L*^*^. An alternate manner to describe self-similarity is via the forms

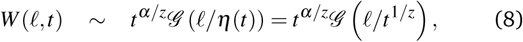

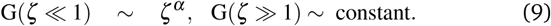

The spatiotemporal evolution of a flat interface occurs as *W* ∼ *t*^*β*^ at short time; eventually correlations develop along the interface that saturate when the correlation length becomes the characteristic length. The set of exponents then decides the specific universality class describing the growth ^42^. We note that equations (7)- (9) usually describe process without an imposed external length scale. The Edwards-Wilkinson (EW) class describes an interface that roughens due to random particle deposition and is smoothed by lateral motion, such as at the surface of settled granular aggregates under gravity. The Kardar-Parisi-Zhang (KPZ) class describes interfacial growth due to random particle depositions and explicitly includes a nonlinear term for growth perpendicular to the interface. Both the KPZ and EW describe the interface through local growth mechanisms.

In order to test if the roughness profiles we obtained from experiments were self-similar, we attempted to plot these in a manner consistent with equations (7)-(9). Figure 4 summarizes these results. We observe in 4(a) that the interface roughness profiles do suggest a self-similar regime when rescaled in a manner consistent with equation (8). Comparing subfigures 4a(i) with 4a(ii) and 4a(iii) further suggests differences with the EW and KPZ type scalings observed in passive systems. In Figure 4(b) we see that scaled roughness profiles from 4 experiments show similar qualitative features. We see domains where the rescaled curves collapse suggesting a self-similar form. We hypothesize that variations between experiments arise due to density and activity changes; these in turn impact the size of the collective features that interact with the interphase region.

**Fig. 4.**
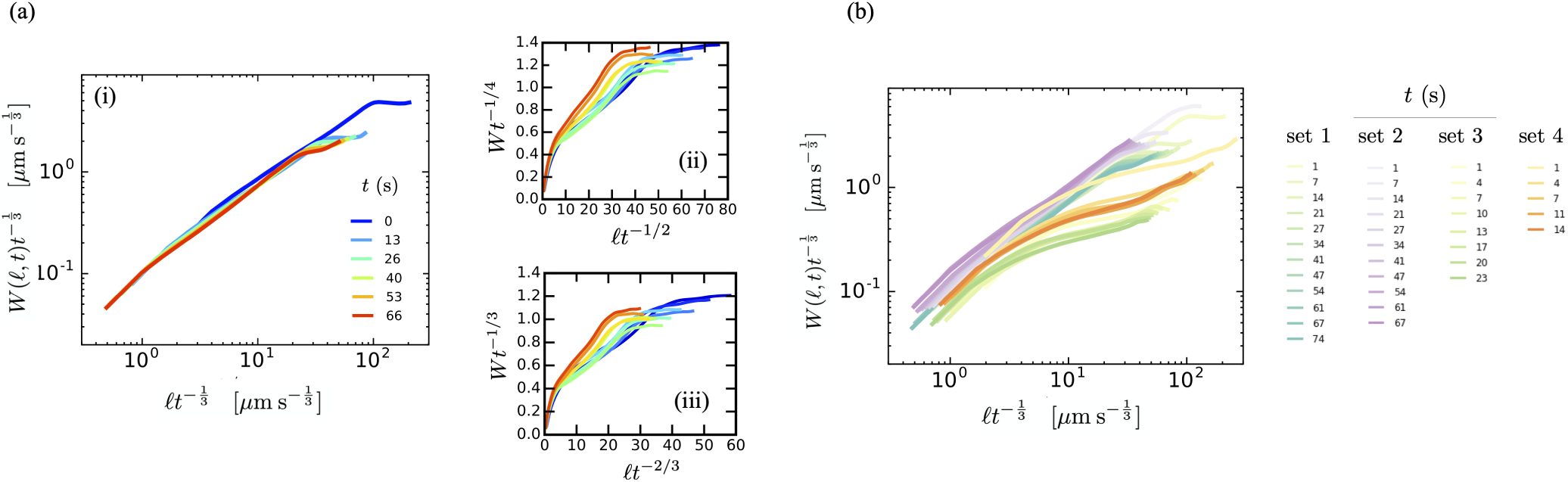
We estimate the roughness and dynamic exponents by measuring the extrinsic width *W*(*ℓ, t*) over time. (a) (i) To determine which scaling exponents best match our system, we try to collapse our curves. Rescaled data points from from Fig. 3a collapse for growth exponent *b* = 1/3 and dynamic exponent *z* = 3. (Right) Curves collapsed using scalings from (ii) EW and (iii) KPZ models. (b) The scaled interface roughness profiles collapse for four separate experimental realizations with each experiment collapsing into its own curve The growth exponent is *b* = 1*/*3 and the dynamic exponent is *z* = 3. Here, we show four experiments, denoted as set 1-4; each curve is plotted using a different time value in that experimental set.

#### 2.1.7 Summary of experimental observations

In summary, our experimental data suggests that strong collective flows in the interphase region play a critical role in the interface structure and propagation. The passive region comprises of densely packed aligned bacteria arranged in static domains that are typically smaller than the structure of the collective flow structures. Over time scales associated with many lifetimes of these structures, the passive region appears as a highly frictional domain which the active bacteria have to penetrate in order to propagate through the medium. Additionally flow induced advection may transport passive bacteria away from the passive region and into the active region. These modalities of surface reorganization are different. For the former, active bacteria in the form of flocks or streamers penetrate the passive region more efficiently; vortices do not. For the latter (advection of passive bacteria), vortical structures are more efficient while streamers and flocks merely serve to push the passive bacteria aside. In addition, observed collective flows include sterically induced flows wherein bacteria push their neighbors in a coordinated manner as well as hydro-dynamically generated flows where flow fields generated due to a moving bacterium influence its neighbors. Both steric interactions and hydrodynamics can lead to enhanced alignment ^40,41^ which in turn lead to faster propagation speeds of bacteria entering the passive phase.

## 3 Simulations of bacterial swarms

It is of interest to understand if steric effects and hydrodynamic effects play equal roles in the interface propagation and in determining the spatiotemporal features of the interface, specifically roughness features and swarming intensity. Furthermore, understanding the relative importance of hydrodynamic interactions will shed light on swarming in moist rather than wet environments. Here, bacteria swarm in a film sufficient to maintain their swarming phenotype. The film of liquid is however not thick enough to allow for long range hydrodynamic interactions. Our experiments on *Serratia marcescens* do not allow us to discriminate and separate the impact of purely steric effects from fluid mediated hydrodynamic interactions on the collective motion well within the active phase as well as in the vicinity of the interface.

To investigate each effect separately and when acting in tandem, we next propose and analyze discrete agent based simulations that treat the bacteria as inertialess self-propelled rods (SPR) ^28^ moving in a viscous fluid. Steric interactions are implemented in a straightforward manner as explained below using standard interaction potentials that depend on distances between the bacteria with rods interacting with all neighbors within a cut-off distance. Hydrodynamic interactions are implemented by treating each rod as a moving line of force dipoles. We do not include thermal noise in our system and ignore translational and rotational diffusion of the rods.

### 3.1 Agent based simulation model without thermal noise

The simulated bacterial swarm system consists of *N* = 56, 000 rod-like bacteria. Each simulated bacteria is idealized as a slender rod of length *ℓ*_b_ and width *λ*(the aspect ratio is thus *a* = *ℓ*_b_*/λ* = 5) that can self-propel. These *in-silico* bacteria move in a domain of area *A* = 500 × 800 (measured in bacterial width units); the ambient fluid is Newtonian with shear viscosity *µ*, and density *ρ*. The rods have an intrinsic self-propulsion speed *u*_*o*_. We note that for *Serratia marcesens*, the self-propulsion speed *u*_*o*_ ∼ 28*µ*m/s swimming in a water-like Newtonian fluid, the Reynolds number *ℛ ≡ ℓ*_b_*u*_*o*_*ρ*/*µ* ≪ 1 and thus the use of the Stokes equation is appropriate. In accord with the experimental settings, we simulate a dense suspension with an area fraction *ψ* = *Nℓ*_b_*λ/A* = 0.7. In simulations, lengths are scaled by rod width *λ*, velocities are scaled by intrinsic self-propulsion velocity *u*_*o*_, time by *τ* = *λ/u*_*o*_ ∼ 0.04 s, and forces by *µu*_*o*_*λ*. Experiments on bacteria in the dilute pre-swarming phase (Figure 5a) suggest that bacteria in colonies have a range of bacterial lengths with a mean between 5-7 *µ*m; longer bacteria that are 2-3 times the length are possible.

**Fig. 5.**
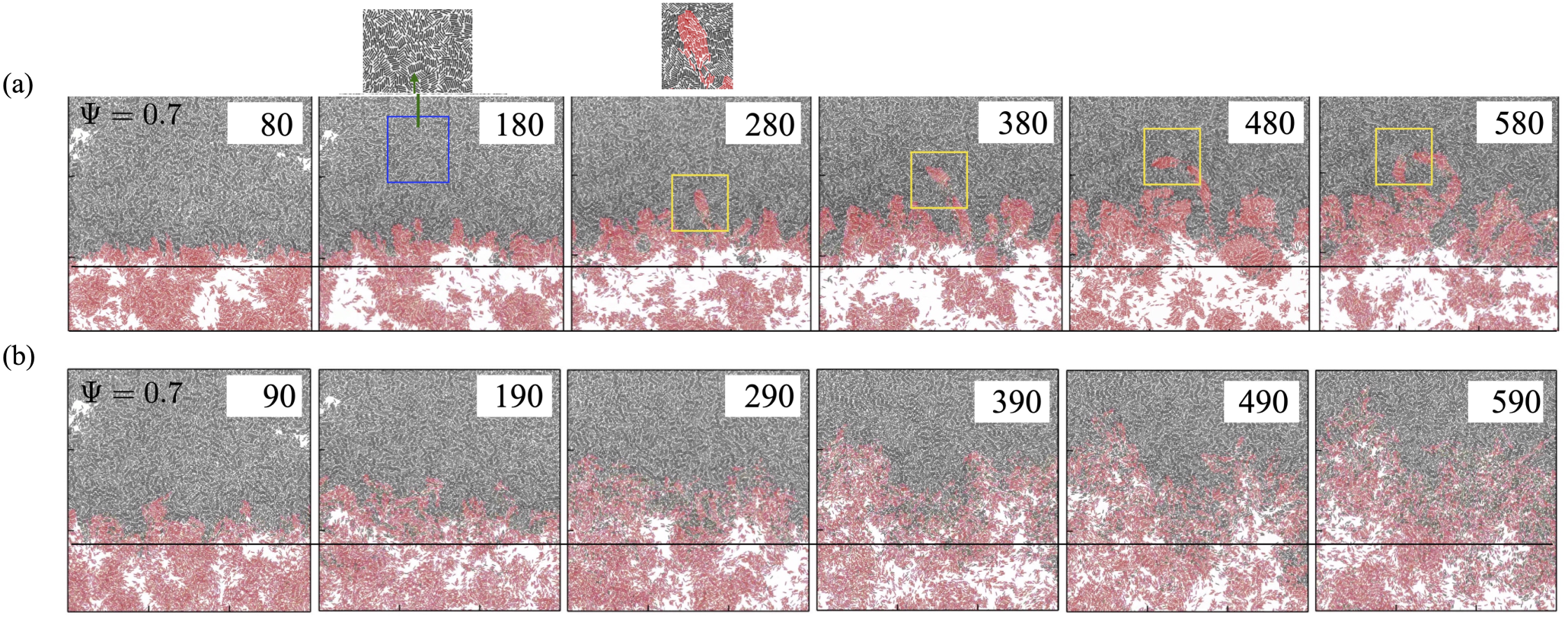
Structural features manifest during the evolution and progression of the front (a) without HI and (b) with HI are highlighted in these snapshots. Detailed time sequences show configurations for active (red) and passive (black) bacteria. The snapshots correspond to ESM Movies 2(a) and 2(b). Here, the simulations use an areal density Ψ = 0.7. The highlighted blue indicates that the passive region in the simulation provides a highly dense, frictional medium with structural features that are small. The initial density is such that the passive phase compacts during the initial stages of the simulation. The highlighted yellow boxes highlight the formation of a finger, that then pinches off and subsequently propagates through this dense frictional passive region. Some such fingers are also observed to break-up into clusters and islands of active bacteria; these penetrate rapidly - the manner by which these intercalate through immotile bacteria is similar to mechanisms observed in our experiments (see also ESM Movie 1, & Figures 1(e) and 1(f)).

Rods are indexed uniquely so as to be able to track them over time. The position and state of a rod with index *α* is completely determined once the position of its mass **r**^*α*^, its orientation vector **p**^*α*^, its linear velocity *d***r**^*α*^ */dt*, and its angular velocity *ω*^*α*^ are known. In the overdamped limit characterized by small Reynolds numbers, the evolution of these variables is governed by

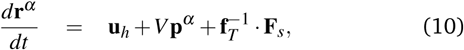

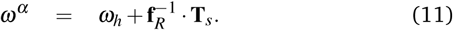

Here, **u**_*h*_ and *ω*_*h*_ are the hydrodynamic contributions to the translational and rotational speed of the rod, respectively *V* = 1 is the dimensionless self-propulsion speed, **p** is the unit vector of rod orientation, **F**_*s*_ and **T**_*s*_ are the force and torque due to the steric effects between neighboring bacteria. The tensors **f**_*T*_ and **f**_*R*_ in (12) and (13) are the translational and rotational friction tensors given by

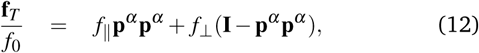

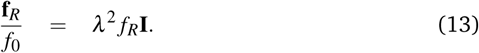

with *f*_0_ = 1 being the Stokesian friction coefficient proportional to the fluid viscosity in scaled units. The three scalar friction coefficients *f*_*∥*_, *f*_*⊥*_ and *f*_*R*_ are dimensionless geometric factors that depend on the aspect ratio *a* of the rods. Here, we adopt the expression for cylinders and use

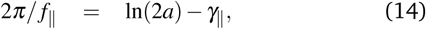

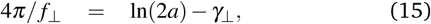

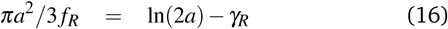

with (*γ*_*∥*_, *γ*_*⊥*_, *γ*_*R*_) given by Broersma ^49^.

### 3.2 Steric interactions

Steric interactions between rods (bacteria) are treated in a discrete manner. We partition each rod into *n* = 6 segments with each segment represented by a point. The position of these points may then be used to evaluate interactions. Geometry dictates that two rods *α* and *β* interact only if their center-of-mass distance |Δ**r**^*αβ*^ | = |**r**^*α*^ − **r**^*β*^ | is less than *ℓ*_b_ + 2*λ*. Let 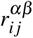 be the distance between the *i*-th segment of rod *α* and *j*-th segment of rod *β* calculated as

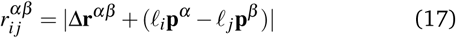

where **p**^*α*^ and **p**^*β*^ are the orientation vectors of rods with indices *α* and *β* and parameters *ℓ*_*i*_, *ℓ*_*j*_ ∈ [−*ℓ*_b_*/*2,*ℓ*_b_*/*2] locate positions of the segments along the rods. The total steric derived potential evaluated along rod *α* is then obtained using the Yukawa potential for the interaction ^28,50^:

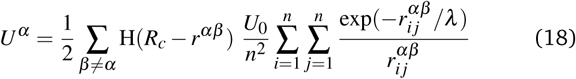

where H(*x*) is the Heaviside step function, *U*_0_ = 800 is the dimensionless potential strength with scale of *µu*_0_*λ* ^2^, and *R*_*c*_ is the cut-off radius beyond which the steric interaction is zero. Here we choose *R*_*c*_ = 1.25*λ*. A linked-cells algorithm for short-range interaction ^51^ is used to accelerate the calculation of the steric interaction given the large number of rods in our simulated system.

### 3.3 Hydrodynamic interactions

The hydrodynamic terms **u**_*h*_ and *ω*_*h*_ are obtained by solving for the background flow experienced by rod *α* due to the dipolar flows generated by all the rods in the system. This background flow is obtained by solving the appropriate Stokes equation in 2D - here, the two-dimensional incompressible Stokes equation for the fluid velocity field **u** supplemented with extra source terms corresponding to the distribution of stresslets (dipolar terms) along each rod in the system. The resulting extra fluid induced drag forces are calculated using the distributed Lagrange multiplier method ^52,53^ used earlier in suspensions of active swimmers.

The flow field induced in the fluid due to the distributed active stresslet is given by

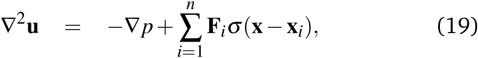

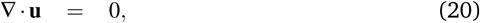

where *p* is the pressure, and **F**_*i*_ = (2*α*_*h*_*/n*)(*ℓ*_*i*_*/*|*ℓ*_*i*_|)*V* **p** is the force distribution for each rod segment, where *i* is the index of rod segments, and *α*_*h*_ is the dimensionless stresslet strength of the bacteria. In our simulations, *α*_*h*_ = 1^21^. In (19), the term *σ*(**x** − **x**_*i*_) converts the forces on rods (Lagrangian grid) to the background flow field (Eulerian grid) with **x, x**_*i*_ denoting the position of Cartesian grid points and rod segments, respectively. The resulting flow field **u** is then interpolated on the rod segments as **v**, and then integrated along the rod length to obtain the total hydrodynamic contributions to the translational and rotational speed of the rod

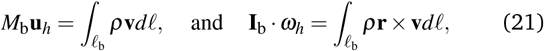

where *ρ* = 1 is the density ratio between rods and the fluid, *ℓ* is the distance along the rod length, **r** is the position vector evaluated from the center of mass, and *M*_b_ and **I**_b_ are the dimensionless mass and moment of inertia of the rod, respectively.

Evaluation of **u**_*h*_ and *ω*_*h*_ closes the set of equations (10)-(16) for the system of rods. Positions and orientations can be tracked in time. Concomitantly, (19)-(21) provides spatiotemporal features of the fluid.

## 4 Simulation results and analysis

### 4.1 Initialization

In the experiments, the quenched passive bacteria in the exposed region were observed to have significant local nematic alignment ^25^. However on the whole when averaged over length scales much larger than 20 *µ*m alignment was more or less random. This is consistent with the paralyzed retaining order at length scales comparable to the size of the collective structures in the active region. The swarming bacteria, when spatially and temporally averaged, are on average randomly oriented (ignoring the small bias due to the speed of the swarming front). Therefore, to maintain consistency across simulations, and to try to mimic this feature in our noise-less simulations, we conducted each simulation in two steps with the domain held fixed. To reduce the uncertainties, we averaged the roughness results based on ten simulation runs with each simulation starting from a different initial rod configuration.

The first step of the simulation allowed us to generate random orientational and spatial distribution of bacteria (rods) at high volume fractions. In order to achieve this, we set all rods to be motile and hydrodynamically interacting; the rods are located initially in a rectangular grid and are aligned. A perturbation is then superimposed on the rod orientation **p**. We let each rod move and evolve for a long enough time so that the position and orientation of the rods represent a possible configuration for the bacterial system before the exposure to light. There are nematic structures in the initial state of the rods, and some regions are void of bacteria because of the swarming effects. It is likely that in the swarming bacteria experiments, noise and possible two-dimensional effects where bacteria can move in and out of the plane may reduce the propensity for such regions devoid of bacteria. The rod positions and orientations are then extracted and used as the initial configuration in the second step.

In the second step, only rods located in the lower half of the domain are set to be motile (*V* = 1, Equation (10)) while the rest are passive (*V* = 0, Equation (10)). We then advance time with each rod moving and interacting with other rods. As the simulation proceeds, we track the active-passive interphase. Here, the simulation domain is a rectangle of a fixed height *H* = 800 with wall boundary conditions applied at the upper and lower boundaries. This height is large enough that the active-passive interphase is not affected by the confinement of the walls initially over the time-scale of our simulations. Periodic boundary conditions are applied to the left and right boundaries of the simulation domain.

### 4.2 Defining the effective interface position

To calculate the number density and velocity fields of bacteria, we divided the simulation domain into small bins of size *δ* = 4 and computed the numbers of both active and passive rods in each bin. We smoothed the velocity and vorticity fields by analyzing the data in overlapping cells ^28^, with cell size 1.5 times bin size.

In our experiments, individual bacteria could not always be clearly visualized especially in the passive region - however their motion was clearly observable. Therefore, we used intensity fluctuations (that are correlated with density fluctuations) to estimate the phase-field order parameter given by equation (3). *Note that fluctuations in intensity are tracked not the intensity itself*. The filtered phase field has value −1 corresponding to the completely passive phase and +1 corresponding to the fully active phase. The interface location and shape obtained thus was consistent with the interface obtained using velocity fluctuations ^24,25^.

In our simulations however, rods can be directly tracked and there is no direct analogue to intensity fluctuations. We used a modified order parameter based on number densities (the subscript S denoting simulation)

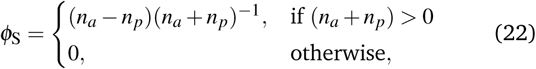

where *n*_*a*_ and *n*_*d*_ are the number density of rod segments for active and passive bacteria, respectively. Here *ϕ*_S_ = 1 corresponds to the purely active phase and *ϕ*_S_ = −1 corresponds to the purely passive phase. The exact value *ϕ*_S_ = 0 corresponds to the regions of equal number of active and passive rods, or the regions that are void of bacteria. We focus attention on the first case; the second where voids can form in the active phase is interesting in itself and is connected to large scale density fluctuations observed in active matter models ^23,32^.

Note that equation (22) exhibits a feature that is not observed or cannot be discerned from experiments. The rod simulations sometimes show voids where there are no rods (active or passive) and hence by definition *ϕ*_*S*_ = 0 since *n*_*a*_ + *n*_*p*_ = 0. This should be distinguished from the case where *n*_*a*_ = *n*_*p*_. Consequently, we will distinguish the manner in which the phase-field is evaluated to zero in subsequent discussions.

Motivated by the success of the phase-field approach in tracking the experimentally observed interphase location, we employ a similar method. The interface height *h*, is calculated by fitting *ϕ*_S_ in a vertical slice with *ϕ*_S_ = tanh((*y*−*h*)*/d*). This calculation of interface is capable of dealing with possible overhangs and mixing in the active-passive interphase. The interface roughness is subsequently evaluated by using the same equation as in the experiments - equation (5).

### 4.3 The propagating interphase region - phenomenology and structure

We first investigated the rod configuration and averaged density and velocity/vorticity fields in the active-passive interphase for self-propelled rods in two limits - without hydrodynamic interactions between rods (no HI), and with full Stokesian hydrodynamics (with HI). Figures 5 and 6 illustrate striking features of the interface motion and suggest significant changes in interface structure when HI is present.

**Fig. 6.**
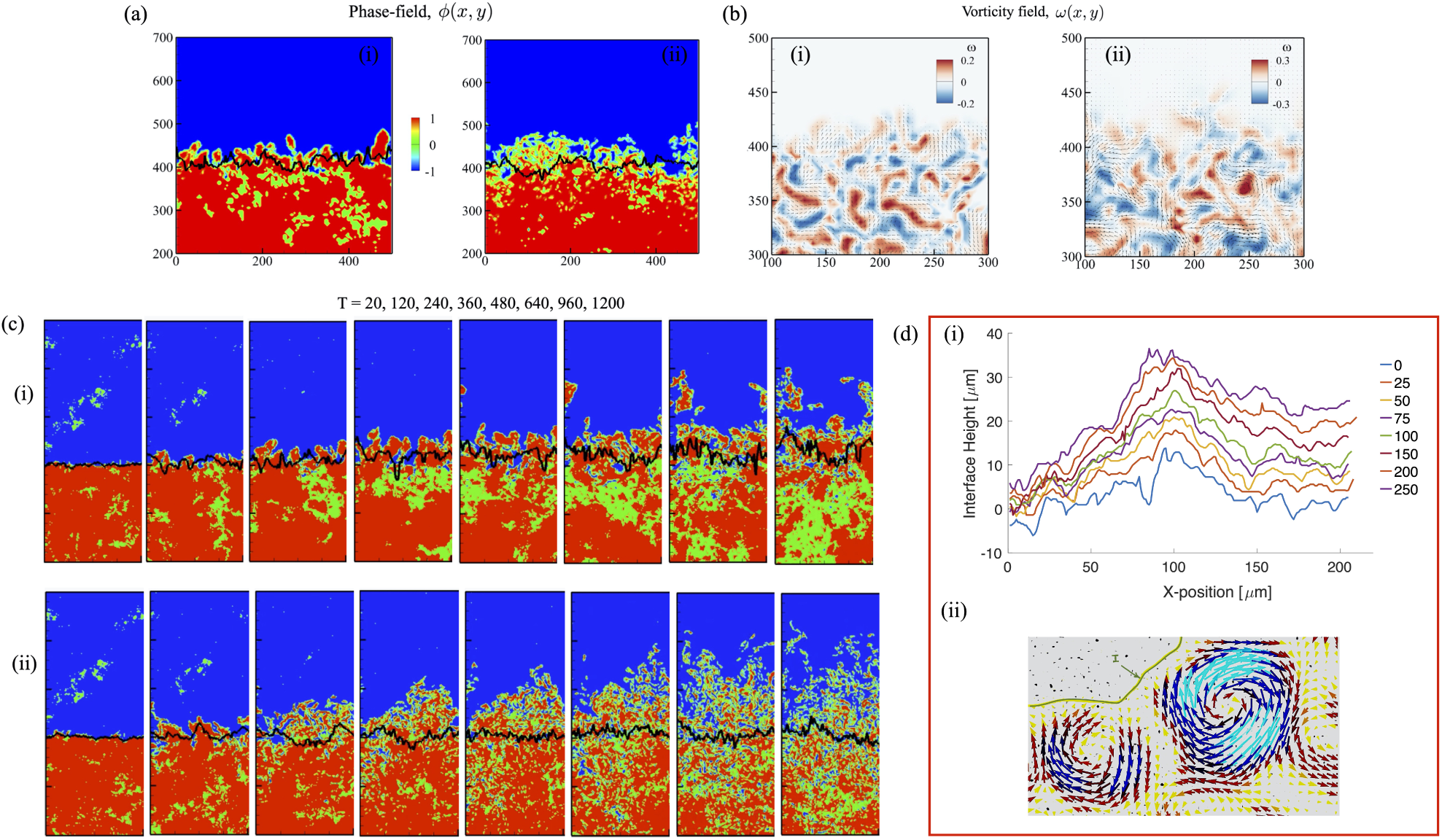
(a) Order parameter scalar fields *ϕ*_S_ for simulations depicted in Figure 5 defined using equation (22) with HI (left) and without HI (right). The full time evolution may be observed in ESM Movies 3a and 3b. Here, *ϕ*_S_ = −1 represents the purely active phase, and *ϕ*_S_ = 1 represents the purely passive phase. Here, *ϕ*_S_ = 0 can represent regions of no bacteria or characterize equal density of active and passive bacteria. The interface between active and passive phases is marked using the black curve. Black irregular curves represent the calculated position of the interface extracted using the hyperbolic tangent function. Note that strictly speaking this approach is valid only under quasi-static conditions i.e, when interphase evolves very slowly and in a manner where the motion is averaged over many time-scales of the local active flow structures. (b) The simulated vorticity field of the swarming bacteria for both cases illustrating length scales of the spatiotemporal features. ESM Movies 4a (without HI) and 4b (with HI) provide more details of the time evolution and persistence time of these features. We observe significant regions with vortical structures. (c) Here, we highlight the role of hydrodynamic interactions in significantly modifying both the interface morphology, effective thickness as well as propagation speed. We note that the interface without HI (set of tiles labelled (i)), is less diffuse than the interface when hydrodynamic interactions are present. We note also the increase in voids in the active phase, fast moving large-scale flocs traveling in the passive phase and the significantly lower mixing in the absence of HI. Finally comparison of the jagged black lines - the computed interface heights and profiles - suggests that the effective roughness is also dependent on the presence of hydrodynamic interactions. (d) (i) Experimental results for *h*(*x, t*) (time stamp here in seconds), illustrating an instance where a local protuberances first forms and then grows (forming fingers) and intrudes into the passive region. However the fingers are not as pronounced; suggesting a response intermediate between the dry state (no HI) and fully wet state (HI dominates). (ii) These sharp features of the evolving interface are typically connected to the presence of sharp vortical structures (either rotating the same way or in opposite senses) that stretch and convect material points (bacteria) in the interface region. The position of the interface is marked in green.

When the hydrodynamic effects are excluded (Fig. 5(a)), the rods only interact with each other through the short-range steric interactions. In the active phase, bacteria have a pronounced tendency to form collective structures such as flocks and streamers; these are nematic structures with groups of bacteria are moving towards the same direction. The bacteria comprising these structures are more likely to penetrate and form incisive cracks in the passive phase. Eventually these protrusions form finger-like or island-like (if these are disconnected from the active phase) regions in the passive phase. Since the passive bacteria cannot enter the active phase due to being immotile and have to be convected by the incoming active bacteria, there is net flux of bacteria from the active phase to the passive phase. Right below the active-passive interface, there we observe significant number of voids where very few active bacteria are present. Furthermore, voids span multiple bacterial lengths.

In summary, the active-passive interface is always moving towards the passive phase; trapping of the active bacteria in the passive domain causes the bacterial number density in the passive phase to increase. We note that the motion of these flocks and streamers bears resemblance to the manner in which individual bacteria align, form clusters and then push through immobile or static bacteria (Figure 5a, and ESM Movie 1).

When hydrodynamic interactions are included as shown in Figure 5(b), active and passive phases are better mixed and the fingering seen previously is not observed possibly due to HI induced destabilization. The length scale of nematic structures generated in the passive domain decreases due to the hydrodynamically generated active flows. Passive bacteria are displaced and advected through the active region in both cases eventually leading to mixing. We anticipate that continuous realignment and instabilities are due to local shearing flows generated over the scale of many bacteria. Note again that we do not have diffusion in our simulations, and hence this destabilization is from hydrodynamic effects; the final result is an active, turbulent motion of rods ^21^ with highly vortical structures. The flow vortices lead to breakup of the active-passive interface and substantially enhance the mixing between the active and passive phases.

Comparison of sequences Figure 5(a) and Figure 5(b) (also ESM Movies 2a,2b and 3a,3b) clarify the nature of structural differences in the interface region, and the variations in the manner in which the leading front propagates without and with HI. Hydrodynamic interactions are associated with a reduced occurrence of dendritic protrusions that may pinch off, penetrate and move through the frictional passive region. Furthermore, we see larger (white) voids with lower values of local densities when hydrodynamic interactions are neglected. We also note larger local interface mixing speeds when hydrodynamic interactions are included. The interphase region is more smeared out and exhibits the hallmark of a mushy, bi-phasic mixed region when hydrodynamic interactions are present.

These conclusions are clarified and supported when the computational results are recast in terms of the density dependent phase field *ϕ*_*s*_ (see also ESM Movies 3a and 3b). The phase field calculated using equation (22) is shown in Figure 6(a), and the interface profiles are indicated on the same figure as the black curve. In both cases - with and without HI, the interfaces start from a flat surface and then roughen.

Figure 6(a) is a representative snapshot of the structure of the phase-field during the front evolution focussing on the interphase morphology and Figure 6(b) shows the associated calculated vorticity fields (note that 6(b) is a sub-area of the region shown in 6(a)), again focusing on the interphase region. As anticipated from Figures 5(a) and 5(b), we observed the following. (i) The front in the absence of HI clearly demonstrates fingering and clustering with these pinching off and moving deep into the passive frictional domain. (ii) Non-HI simulations demonstrate significant density variations in the active phase, reminiscent of the large amplitude density fluctuations seen in model theoretical active polar and nematic systems ^21,28^. These fluctuations are usually smeared out when hydrodynamic interactions are allowed. (iii) Finally, again as expected, HI smears out the interphase region which now possesses features of a diffuse interface. Note that a finite interface thickness was calculated in the *Serratia* experiments and reported earlier by us ^24^ using both intensity variations (a stand-in for density variations) as also independently using PIV. Here we see that hydrodynamic interactions significantly thicken the apparent interface - i.e, there is truly a well mixed bi-phasic region when HI is present.

Examination of the associated vorticity fields shows that vortical structures are slightly larger and more segregated when hydrodynamic interactions are non-existent. Comparing figures 6(b)-(ii) and 6(b)-(i) we conclude that the magnitude of vorticity attained in the interphase region are the same order. It is likely however that the spectral characteristics are different. We will revisit the vorticity field later in this article in following sections. For rods without hydrodynamics, the vortices are elongated and connected due to the presence of the nematic structures. When hydrodynamics are included, the vortices are broken, creating meso-scale turbulence and leading to a more disordered flow field. Here, the magnitude of flow velocity and vorticity are larger than those without hydrodynamics interactions, and there is a slight flow in the passive phase since the hydrodynamic interaction decays slowly with distance.

Hydrodynamic interactions impact not just the morphology and structure of the interphase region but also the evolution and dynamics. The striking difference between simulations without and with HI are evident in Figures 6(c)-(i) and 6(c)-(ii). Note the increasingly bi-phasic structure of the interphase region due to hydrodynamic interactions; additionally, this mushy region gradually increases in width due to mixing. Green domains deep in the active phase observed in Figure 6(c)-(i) corresponding to voids or low density regions are significantly less for the case with HI than without.

### 4.4 Vorticity fields and velocity correlations

To check the effect of the bounding walls as well as to investigate how velocity fields observed in our simulations compared with experiments, we next analyzed the vorticity distribution and velocity correlations far from the active-passive interphase region. Figure 7 and ESM Movies 4a and 4b provide a visual representation of our results. Figure 7(a) shows the vorticity distribution as well as the location of the boundary between the phases near the beginning of the simulation (*T* = 20, left) and then well into the simulation (*T* = 220, right). Here hydrodynamic interactions are turned on. We note that vorticity magnitudes across the active region are consistently similar once we move away from the interface and deep into the active region (yellow box). As time progresses however the distance between vortical structures increases slightly possibly due to the mixing and convective flux of passive rods into the active phase.

**Fig. 7.**
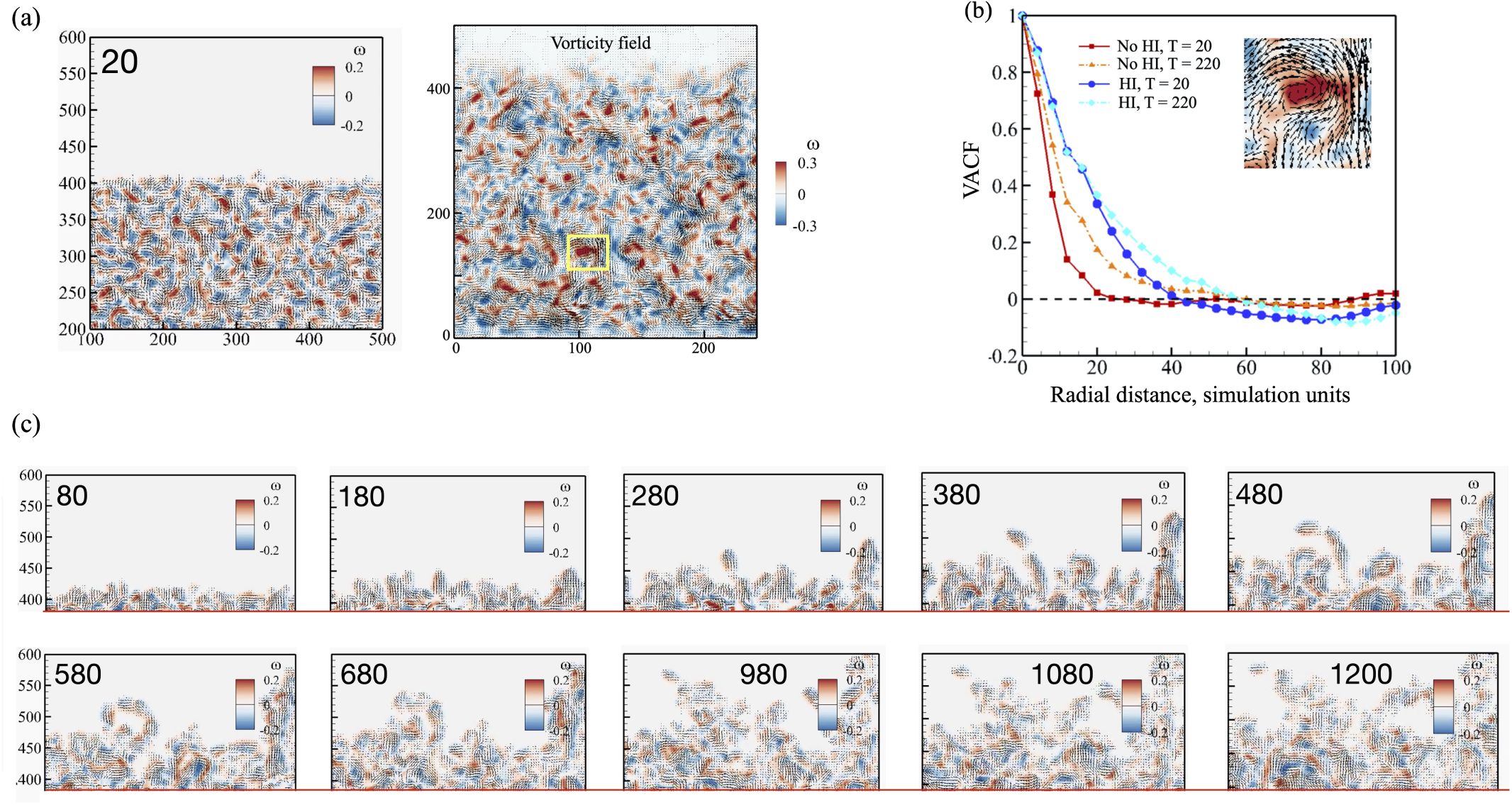
(a) (Left) (Right) The vorticity field in active phase is shown in more detail here illustrating interspersed clockwise and counterclockwise vortical features and large-scale streaming flows. (b) Velocity correlation function (VCF) calculated in the active phase are plotted here for two time instants and for cases without and with HI. The characteristic vortex radius is the radius at which VCF=0. (Inset) Close up of the vortical structure in the yellow box indicated in ((a) - right). (c) Close up snapshots of the vorticity field at the interfacial region illustrating salient morphological features for dry simulations (without HI).Time stamps are indicated as insets in (a) & (c).

A closer examination shows wall effects on the active phase for roughly *y* < 50; the no-flux condition causes rods to reorient and move parallel to the wall leading to a velocity in the same direction and negligible vorticity. In the bulk of active phase (∼ when 50 < *y* < 350), clear vortical structures exist (Figure (7(a) right). Vortices of both signs form periodically in a manner vey similar to experiments ^24^.

To quantify the structural features of the velocity field, in addition to the vorticity field, of the vortical structures in this region, we calculate the velocity correlation function (VCF) in Figure 7(b) choosing the region 100 < *y* < 300. Furthermore, to discern effects due to HI, we analyze simulations with and without HI. Given the VCF, characteristic vortex sizes can be found from the length-scale at which VCF equals 0 and crosses over to positive values. The results at *T* = 20 correspond to pure active phase since the density of passive rods in the active phase is negligible. At the start of simulations (*t* = 20), the vortex radius for bacteria without HI is about 25*µ*m and for bacteria with HI is 40 *µ*m. The vortex radius is 40 *µ*m for a dense suspension of *B. subtilis* ^20^, *and is about 20 –* 25 *µ*m for swarming *Serratia marcescens*. Also at *T* = 20, we observe that velocity for the rods without HI (red curve, red filled circles in the figure) is less correlated than rods with HI (blue curve, blue filled circles line in the figure). The vortex size for active bacteria phase is about 40 microns (agreeing with our experiments). As time goes by, for either with or without hydrodynamics, the number density of active bacteria at the studied region decreases because some active bacteria penetrate into the passive domain.

Associated with the decrease in number density is an increase in vortex radius. Interestingly, these conclusions are consistent with our reported observations on slightly increases in size and longer residence time of vortical structures close to the inter-phase ^24^ (and thus higher density of passive rods). Thus our computations capture qualitative features of the swarm and predict correlation lengths consistent with experiments.

### 4.5 Roughness of the interface and comparison with experiments

Finally, combining experiments with simulations, we conclude that hydrodynamic and steric interactions enable different modes of surface dynamics, morphology and thus front invasion. This is clearly seen when one compares the vorticity fields in the inter-phase region for simulations with and without HI (Figures 5(a,b), Figure 6 and Figure 7(c)). In the absence of HI such as for dry systems where the only contribution from ambient fluid is the Stokes drag on the rods, we observe a sharper interface structure, albeit with a more disruptive morphology and extensive fingering. Flocks and streamers can develop at the interface, and exploiting any structural weakening in the passive phase can pinch off and penetrate the frictional medium. What is noteworthy is that these clusters are also characterized by strong vorticity and thus involve circulatory motions.

Comparing the jagged structure of the interface as indicated by the black curve in Figures 6(c)-(i) and 6(c)-(ii), we hypothesize the different modalities of surface deformation and evolution not only result in different propagation speeds and effective interface thicknesses but also lead to different roughness. Figure 8 confirms this hypothesis - note that in obtaining the roughness profiles, we varied the width of our simulation domain, which is denoted as L.

**Fig. 8.**
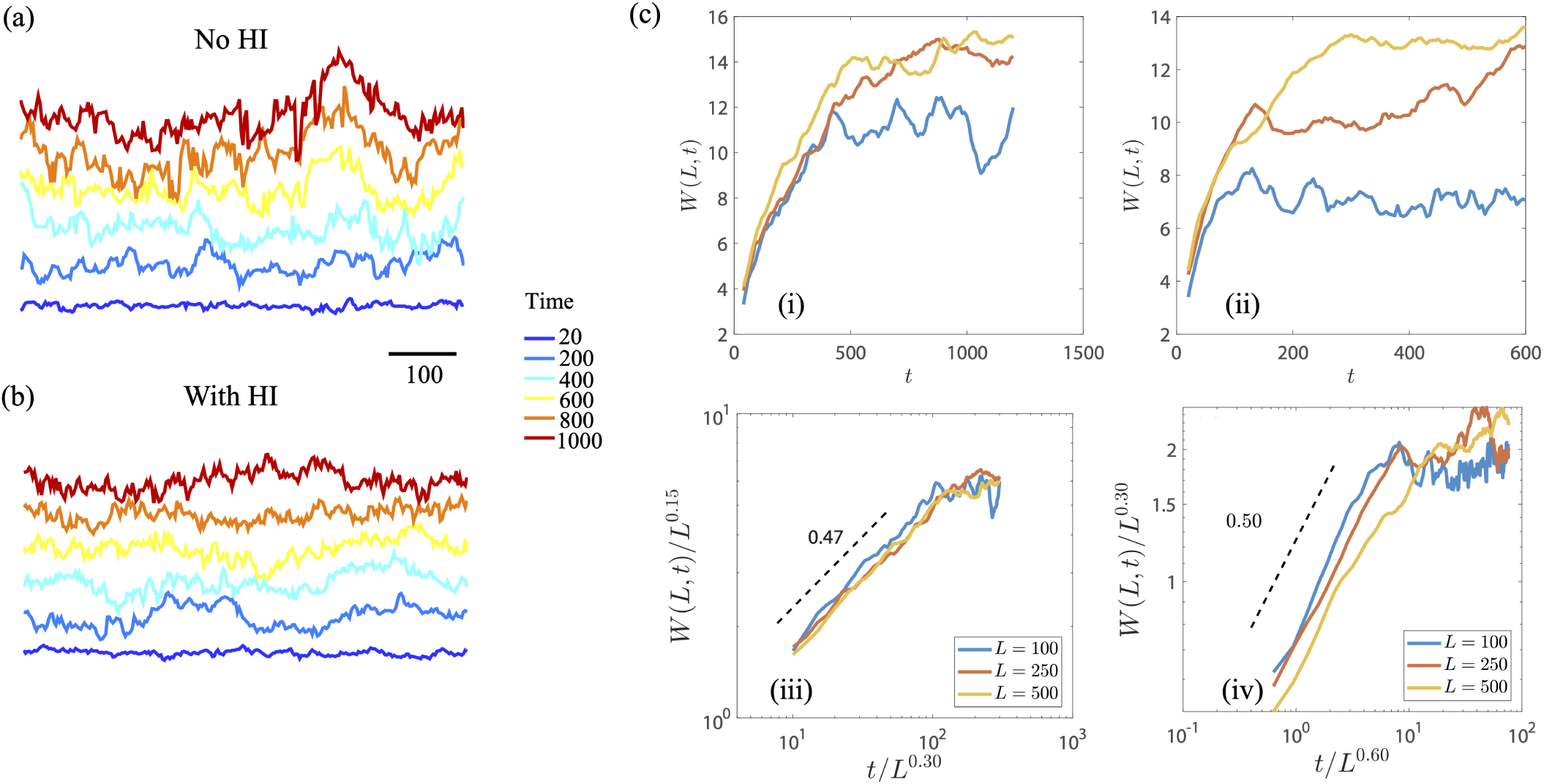
Interface roughness *W*(*L, t*) evaluated over the whole system as a function of the time for different system sizes (a,b) Interface profiles in the (*x, y*) plane obtained from the phase-field. For better visibility, profiles at different time are separated by adding a vertical offset. (c) The averaged interface roughness scaled by the system size as a function of scaled time collapses onto a single curve. In (i,iii), hydrodynamic effects are neglected. In (ii,iv), hydrodynamic effects are included.

We next investigated the scaling laws associated with the roughness features, given by *W*, as defined by equation 5, of the active-passive interface. For bacteria either without - figures 8(c) - (i,iii) - or with hydrodynamics - figures 8(c) - (ii,iv) - the interface roughness always increases at early stages and reaches a plateau with discernible saturation times *t*_×_. Motivated by the collapse observed in the experiments and illustrated in figure 4, we then tested if our simulation data could be collapsed onto a master curve by scaling the roughness with *L*^*α*^ and time by *L*^*z*^, where *α* is the roughness exponent and *z* is the dynamic exponent. The results are shown in Figure 8(c) - (i,iii) for simulations without hydrodynamic interactions and Figure 8(c) - (ii,iv) for simulations with hydrodynamic interactions. The increasing rate in the early stage corresponds to the growth exponent *β*, which is around 0.5 for bacteria either with or without hydrodynamics. The inclusion of hydrodynamics increases *α* and *z*, meaning that the saturation time and the maximum roughness both increase with the system size with larger exponents. In Fig. 8(c) - (iv), the curves do not collapse, probably because the interface is not self-affine.

We also follow the same method as in the experiments and plot the local roughness as a function of box sizes. The interface roughness here is evaluated at simulation time from 600 to 1200, corresponding to 21 s to 43 s in experiments. For bacteria either with or without hydrodynamics, we find that the local roughness increases with the box size first and then slightly increases when the box size exceeds a critical length. When the hydrodynamic effects are neglected, the data can be collapsed by scaling both the roughness and box size by *t*^0.2^. For simulations with hydro-dynamics, this scaling increases to *t*^0.3^, and agrees with those in the experimental results. This is after the saturation time in the simulation (*t*_×_ ∼ 250) so that the exponents are different from the scaling in the roughness growth.

## 5 Summary and Outlook

Swarming is a complex multi-cellular response featuring fast, collective and long ranged intense bacteria laden flows ^5,6^ and enables efficient and rapid colonization of environments and biologically important surfaces. This phenomenon requires the integration of multiple and multimodal chemical and physical signals, including physiological and morphological differentiation of the bacteria into swarmer cells, the optimal presence of nutrient and favorable porous surfaces ^5–7^. A significant component of the swarming state are interfaces or propagating fronts that form naturally allowing swarms to invade free space or highly frictional domains. Here, we study the evolving structural features of an active-passive interfaces using dense colonies of *Serratia marcesens* as a model system. We image the evolution, dynamics and morphology of the interphase region between an active (swarming) phase and a passive (immotile) phase. The passive phase is densely packed and micro-structured and hence offers an highly frictional environment. Integrating results from Particle Image Velocimetry (PIV), intensity based image analysis and analysis of surface fluctuations, we quantify the dynamics and morphology of the evolving interface. We show that correlations between spatially separated surface fluctuations and damping of the same are influenced by the interaction of the interfacial region with adjacently located collective flows. Furthermore, dynamical growth exponents characterizing the active manner in which the interfacial region is continuously reshaped differ significantly from classically expected values.

To complement and interpret our experiments, we presented a high-performance numerical simulation that explicitly tracks the motion of each bacteria with consideration of both cell-cell steric interaction and far-field hydrodynamic interactions. The simulations are able to separate the steric and hydrodynamic effects and illustrate the time evolution of mixing between the active and dead bacteria swarms for either with or without hydrodynamic interactions. The hydrodynamic effects have a tendency to create turbulence in active phase, which disturbs the active-passive interface and thus enhance the mixing between passive and active bacteria. The *active turbulence* also decreases the length scale of the nematic structure and make it easier for active bacteria to penetrate the passive phase. At the same time, simulations in the dry regime - where hydrodynamic interactions are neglected - strongly indicate significant fingering instabilities with macro- scale flocks penetrating the passive phase, pinching off from the rest of the active domain and covering significant territory in the passive region before fragmenting. The interface with hydrodynamic effects shows larger roughness exponent and dynamic exponent than those without hydrodynamic effects, but the growth exponents are similar.

Combining experiments with simulations, we conclude that hydrodynamic and steric interactions enable strikingly different modes of surface dynamics, morphology and thus front invasion. Swarms typically utilize and move in nutrient-laden ambient fluid in a swarm that is extracted from underlying soft poroelastic fluid infiltrated substrates. Our observations imply that surfaces tailored with a friction gradient or a porosity gradient may hinder swarm propagation significantly.

We conclude with some possible extensions of this work. Our results strongly motivate the need for further careful experimentation where bacterial swarm intensity (density and/or self-propulsion), the substrate friction and softness can all be varied independently. This will allow us to investigate carefully the physicochemical and biophysical conditions under which the front propagation and attendant instabilities may be suppressed or enhanced. In turn, this information will also help elucidate the mechanisms underlying the formation of jets and streamers observed in many advancing bacterial interfaces.

The agent-based numerical model we studied considered the propagation of swarms into frictional domains and was motivated by the manner in which bacteria may reconstitute and reclaim territory, and also the manner in which bacteria may physically interact with sub-domains of other bacterial species or mutants with strongly contrasting self-propulsion. It would be of interest to generalize our treatment by allowing the self-propulsion speed and viscous friction to depend on biophysical parameters such as substrate properties. Similarly, introducing a dependence on an ambient diffusing chemical may provide a minimal model of quorum sensing.

Finally, in order to probe the active matter related aspects of swarming matter such as the co-existence and formation of phases, one can combine the agent-based microscopic models presented in this paper with mesoscale models for active liquid crystals ^32–36^.

## Competing interests

Authors have no competing interests.

## Acknowledgements

JT received an undergraduate research fellowship from the NSF-CREST Center for Cellular and Biomolecular Machines and acknowledges support from NSF-HRD-1547848. AMA acknowledges grants from NSF-CBET-1700961 and NSF-CBET-1604423. AG would like to thank Dr. Paulo Arratia for preliminary discussions that lead to this work. Previously published data gathered by AG and AEP and published elsewhere with Dr. Arratia was used to guide the design of the computational agent based scheme.

## Conflicts of interest

There are no conflicts to declare.

## Author Contributions

AEP and AG conceived and performed the experiments. JT, AG and AEP analyzed the experimental data. AMA and AG conceived the simulation model. YZ performed the simulations, and YZ, AMA and AG analyzed simulation results. AG wrote the manuscript along with co-authors.

## Data availability

Relevant data and simulation details are available from AG and AMA. Supplementary movies may also be directly downloaded from https://gopinathlabucm.wixsite.com/cmml/downloads.

## A

### Experiments: Methods and Materials

#### A.1 Sample preparation: Growth of swarms and light exposure

Swarms of *Serratia marcesens*(ATCC 274, Manassas, VA) on agar substrates were prepared by dissolving 1 w% Bacto Tryptone, 0.5 w% yeast extract, 0.5 w% NaCl, and 0.6 w% Bacto Agar in deionized water. Melted agar was poured into petri dishes; to this 2 w% of glucose solution (25 w%) was then added. *Serratia marcescens* was then plated and inoculated on the solidified agar plates and then incubated at 34*o*C. Colonies were observed to form at the inoculation site and then grow outward on the agar substrate. Experiments on these swarming colonies were performed 12-16 hours after inoculation. Figure 1(a) provides a snapshot of the swarm; we observe the leading edge propagating onto free agar (top side, white edge is the interface of the swarm propagating onto free media) while intense swarming is observed towards the lower half of the figure as one moves away from this edge.

To generate an internal boundary separating active swarming domains from a large passive immotile domain, we expose part of the swarm to sustained high intensity light from an unfiltered wide-spectrum mercury lamp with known spectral characteristics. Standard fluorescence microscope optical components were used to focus the light; the light path included a half-plane aperture.

The effects of light exposure on the collective swarming motility of *Serratia marcescens* have been reported previously ^25^. For small exposure times (20 - 40 seconds) and weak intensities (*I* < 220 mW at 535 nm) exposed cells remain active or are only temporarily passivated. For long exposure times (> 60 seconds) and sufficiently high intensities (*I* > 220mW at 535 nm), the bacteria are rendered immotile with large scale motions quenched permanently. For experiments reported here, we chose an exposure time of 60 seconds at intensity *I* = 370 *µ*W (at 535 nm). Intense light under these conditions when projected on the swarm through the half-plane aperture yielded a paralyzed region (passive phase) within a swarming domain (active phase) - the inter- phase boundary between the two was roughly straight at scales larger than bacterial lengths and remained so as long as exposure is maintained. When exposure was stopped, the active phase penetrated the passive phase.

#### A.2 Imaging: Microscopy and PIV tools

High speed imaging coupled with structural analysis based on the *Orientation J* plugin of the open-source software *Image J* ^38^ and Particle Image Velocimetry (PIV) based on the open source software PIVLab ^39^ were used. The open-source software *Image J* and associated plugins were used to analyze the alignment and cluster size of bacteria using static images digitized and thresholded appropriately.

Bacteria were recorded (with the agar plates facing down) using an inverted Nikon microscope Eclipse Ti-U using either a Nikon 10x (NA = 0.3), 20x (NA = 0.45) or 63x (NA = 0.7) objectives. Images were gathered at either 30 frames per seconds with a Sony XCD-SX90 camera or at 60/125 frames per second with a Photron Fastcam SA1.1 camera. We used videos of the swarm and PIVLab software to extract the bacterial swarm velocity fields **v**(**r**, *t*) at locations in the recorded region. The velocity field was sampled at 3 *µ*m spatial intervals, with images taken at 60 or 125 fps. We used algorithms based on direct Fourier transform correlations, multiple passes and window sizes. The initial passes used large interrogation windows for better signal-to-noise ratio and more robust cross correlation. The interrogation window sizes decreased with each pass to increase the vector resolution. The window sizes used were 64, 32, and 8 pixels or 21,11, and 2.5 *µ*m, allowing us to detect bacteria speeds ∼ 50*µ*m/s.

## Footnotes

* These calculate directional derivatives of the gradient in image features, obtain the direction along which these derivatives are maximized, and thereby compute the structure tensor. The structure tensor provides eigenvalues and eigenvectors that characterize the alignment direction, degree of alignment and degree of coherence.

## Notes

### Competing Interest Statement

The authors have declared no competing interest.

### Summary of Updates

Author list modified. Figures and figure captions revised and typos corrected.

